# The Roads Not Taken: The Neural Dynamics of Lexical Alternatives During Free Language Production

**DOI:** 10.1101/2025.11.09.687421

**Authors:** Daria Lioubashevski, Daniel Friedman, Adeen Flinker, Ariel Goldstein

## Abstract

Human language allows multiple ways to express the same thought, raising the possibility that several lexical candidates are activated in parallel before a single word is selected for articulation. While their co-activation is assumed by computational models of word production, until recently, there has been no practical way to estimate the identity of these candidates in unconstrained speech, nor to directly probe their representation in neural activity. Here, we combined high-density electrocorticography recorded during real-world conversation with context-sensitive large language model (LLM) to characterize, for the first time, how these lexical candidates are represented in the brain. We focus on the candidates which remain unsaid, referred to as the “alternatives”, by only using samples where the LLM predictions do not include the produced word. We provide evidence for three computational principles of the neural dynamics of lexical alternatives: (1) multiple lexical alternatives to each word in conversation are distinctly represented in neural activity prior to articulation; (2) the set of lexical alternatives is rapidly updated from one word to the next; and (3) alternatives to future words are represented along with alternatives to the upcoming word. Together, these findings reveal that spontaneous speech production is supported by a broad, dynamically evolving and forward-looking set of lexical alternatives, providing a neural account of how the brain prepares not only the words that are spoken, but also those that remain unsaid.

## Introduction

Let’s say a friend asks you what you thought of their presentation at work. Depending on your opinion, how honest you want to be and how close of a friend they are, you might say it was “amazing”, “great” or just “good”. We know that given multiple options, people usually choose the word that is most aligned with the speaker’s goal in that interaction (Nozari and Novick 2017). This raises the question of whether all of these possible words are represented in your brain prior to articulation, despite the fact that you end up saying only one of them. We call the word that is ultimately said the “target word”, whereas the remaining, unproduced candidates, are referred to as “lexical alternatives”. Studying these alternatives is particularly difficult because, unlike the target word, they are neither articulated nor reliably accessible through introspection. Neural recordings therefore provide a unique opportunity to probe these otherwise covert alternatives as they emerge in real time. Prior evidence for their parallel activation has relied largely on indirect behavioral signatures, such as speech errors (Dell and Reich 1981; Motley and Baars 1976; Rapp and Goldrick 2000) and highly controlled picture-naming paradigms (Spalek et al. 2013), leaving unresolved the question of how their neural representations are organized and unfold during natural language production.

We address this question by identifying three computational principles that characterize the neural dynamics of lexical alternatives: (1) multiple lexical alternatives to each word in conversation are represented simultaneously prior to articulation; (2) the set of lexical alternatives is continuously updated on a word by word basis during an utterance; and (3) alternatives to future words are represented along with alternatives to the current target. We begin by establishing the co-activation of multiple alternatives in real-world conversations (principle 1), and then characterize how they evolve over time (principles 2 & 3).

### Principle 1: Multiple lexical alternatives are activated in real-world language production

Whether multiple lexical candidates are activated simultaneously during real-world language production, and how many, remains an open question even after decades of work on lexical selection (W. J. M. Levelt et al. 1991; Belke et al. 2005; Howard et al. 2006). This is due to several limitations shared by prior studies. First, they provide only indirect behavioral evidence for co-activation, without quantifying how many specific lexical candidates are activated at any given moment ((Schriefers et al. 1990; Hutson and Damian 2014); (Bormann 2011; Fieder et al. 2019)). Second, they rely on highly constrained experimental setups which may not generalize to real-world production settings (Balatsou et al. 2022) (Oppenheim and Nozari 2024). Finally, they offer little to no *neural* evidence for the parallel activation of multiple lexical candidates, in part because speech production is difficult to study with classic imaging methods (Abbasi et al. 2021; Volfart et al. 2024; De Vos et al. 2010).

Two methodological breakthroughs in recent years now make investigation into such questions possible. First, contemporary large-scale electrocorticography (ECoG) datasets capture neural activity with high temporal and spatial resolution during spontaneous natural conversation (e.g., (Goldstein, Wang, et al. 2025; Zada et al. 2024)). Second, large language models (LLMs) have emerged as a powerful new tool for language generation (Radford et al. 2019). LLMs make next-word predictions that closely match those of humans (Goldstein et al. 2022), by generating a probability distribution over the entire lexicon (Fig. 1), essentially ranking all possible words in order from most to least likely. Taken together, these properties mean that the top ranked predictions of LLMs give us for the first time a practical way to estimate, for each word in a real-world conversation, the set of likely lexical candidates. Because our aim is to characterize the representation of (unsaid) alternatives, we restrict ourselves where these predictions do not include the target word. By combining ECoG recordings with LLM-based predictions, we first demonstrate that these alternatives are represented in neural activity prior to target word onset and that each alternative captures a unique portion of the variance in the neural signal (Sections 1.1-1.4). We then estimate the size of the candidate set (including the target word and alternatives), by using increasingly larger sets of lexical candidates to predict neural activity. We find that the explained variance grows with the number of candidates up to twenty words, suggesting more than a dozen words are represented in parallel (Section 1.5).

**Figure 1.**
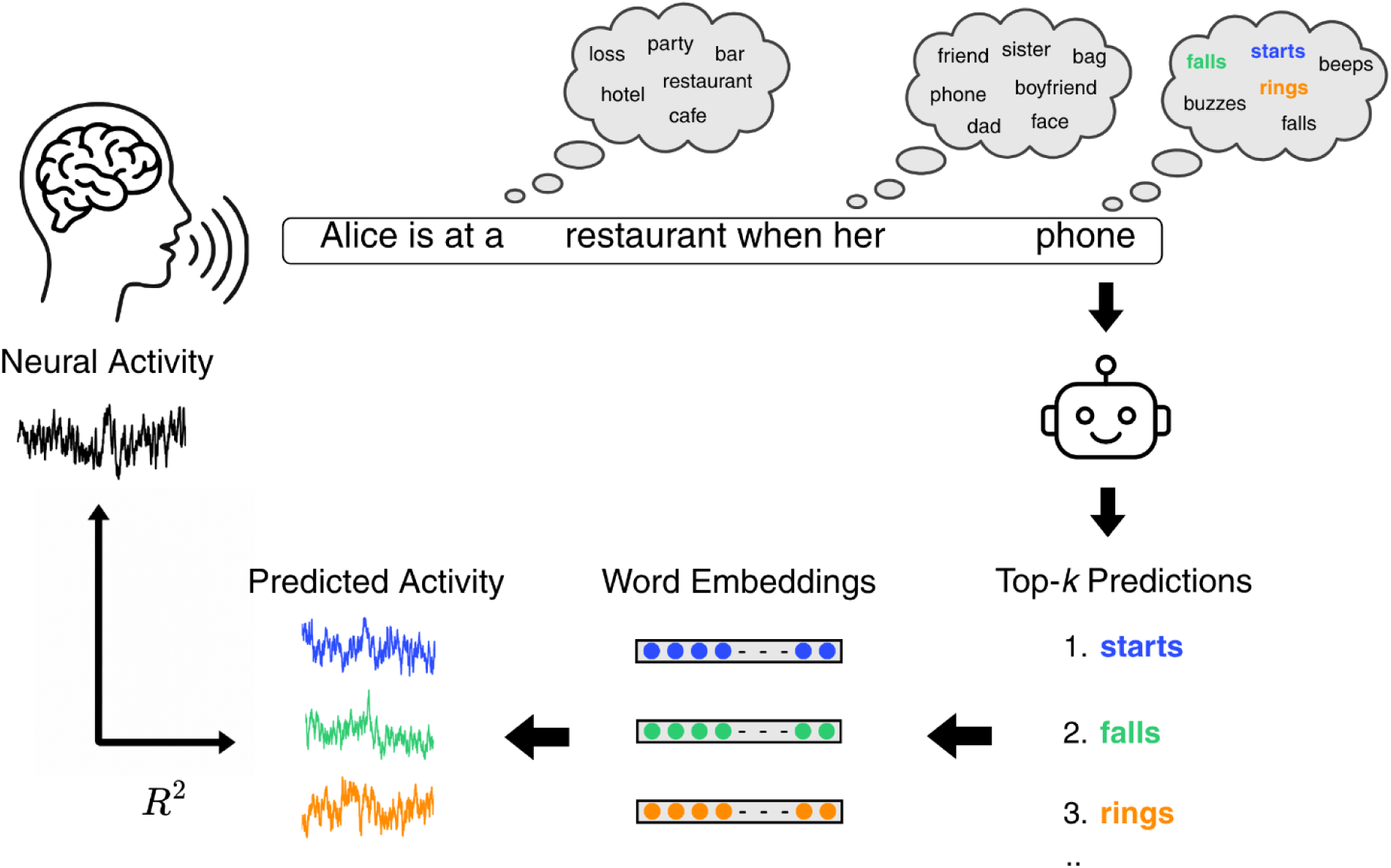
Visualization of our framework for identifying co-activated lexical candidates during real-world production. In the example utterance (“Alice is at a restaurant when her phone…”), thought bubbles illustrate multiple lexical candidates active at different points in the utterance, including the target word and unspoken alternatives. At each point in the utterance, we take the predictions of a Large Language Model (LLM) for the next word (based on the preceding context), and treat them as a ranking over its vocabulary from most to least likely. We then extract the top-3 ranking predictions as an estimate for the set of lexical candidates. Finally, static word embeddings of these candidates are used to train linear encoding models that predict the speaker’s neural activity, allowing us to test whether these words are simultaneously represented in the brain.

### Principle 2: The set of lexical alternatives is updated on a word by word basis

Classic studies have mostly focused on single word production (Dell et al. 2008), and even more recent studies of multi-word production usually only consider short and very structured responses to static visual stimuli (e.g., (Janssen and Caramazza 2011; Jaeger et al. 2012). In contrast, in real-world conversation, speakers have to adapt to a constantly changing context and react quickly to their interlocutor, with the average person speaking at a rate of 2-3 words per second and typically responding within a fraction of a second after their conversational partner finishes (Stivers et al. 2009; Brehm and Meyer 2021). This quick-paced dynamic setting where each produced word changes the semantic, syntactic, and pragmatic context constraining what may be said next, suggesting that the set of lexical alternatives needs to be continuously revised during an utterance. By splitting our conversational data according to word position, we show the brain updates its set of lexical alternatives on a word by word basis (Section 2).

### Principle 3: Alternatives to future target words are activated alongside alternatives to the current target

While the scope of lexical planning in sentence production, that is how many words ahead speakers retrieve as lexical items before they start producing an utterance, has long been debated (Zhao and Yang 2016; Roeser et al. 2019), there is some evidence that speakers can retrieve several upcoming target words before articulation (Lee et al. 2013; Martin et al. 2014; Swets et al. 2014). Given that in models of lexical retrieval, the activation of the target word is accompanied by co-activation of lexical alternatives, this implies that alternatives associated with those future positions are also activated. We demonstrate, for the first time, that before the current word is even spoken the brain already represents unspoken alternatives to multiple future words. Meaning that, prior to articulation of target word *i*, lexical alternatives to target words i+1, *i+2, i+3* are represented in the neural activity (Section 3).

Taken together, our findings provide the first comprehensive picture of lexical alternatives in real-world language production: from demonstrating that multiple lexical alternatives are activated in parallel before target word onset, with each alternative explaining a unique portion of the variance in neural activity; to estimating the number of alternatives, and finally showing that set of activated alternatives updates continuously on a word by word basis, and that it even includes alternatives to multiple future target words.

## Results

To investigate how lexical alternatives are represented during spontaneous speech production, we combined intracranial neural recordings from real-world conversations with word-by-word predictions from a large language model. We used the production portion of the 24/7 dataset (Goldstein, Wang, et al. 2025), consisting of over 500,000 words of spontaneous speech recorded over 100 hours of conversations between four ECoG patients and their conversational partners during the patients’ day-to-week-long stays at the epilepsy unit. Production here refers to all words produced by the patients themselves. Given that we’re interested in the representation of lexical alternatives before the target word is produced, we restricted our analyses to neural activity from -500 to -100 ms relative to target word onset, consistent with estimates of the timing of lexical retrieval relative to articulation (Indefrey 2011; Sahin et al. 2009).

In order to estimate a set of lexical candidates for each word in the conversation we leveraged an LLM (e.g., GPT-2 XL, see Supplementary Material for reproduction with Llama-2) to generate word-by-word predictions using a sliding context window of 1024 tokens over the transcript. For each word, the model produced a probability distribution over its vocabulary, which we treated as a ranked list (Fig. 1), extracting the top-*k* predictions as possible lexical candidates (see Section 1.1 for how a subset of these are identified as unspoken alternatives). In most of our analyses we zoom in on the top-3 predictions specifically, because probability differences among predictions become increasingly small and less reliable at lower ranks (Fan et al., 2018; Holtzman et al., 2019).

### Principle 1: Evidence for the Simultaneous Representation of Multiple Lexical Alternatives

We test the hypothesis that during real-world language production, the brain represents multiple alternatives in parallel at the lexical level, rather than committing to a single candidate at a time. We begin by establishing the representation of multiple alternatives through converging decoding (1.1), and encoding evidence (1.2). We then show that each alternative captures a unique aspect of the neural signal beyond their shared similarity (1.3), and using linguistic feature spaces provide evidence that the representation of each alternative contains both semantic and syntactic information. Finally, we explore how many candidates (including the target word) are represented in parallel (1.4).

#### 1.1 Identity of Lexical Alternatives Can Be Decoded from the Neural Signal

We first asked whether multiple lexical candidates, estimated via top-3 LLM predictions, could be decoded from neural activity preceding word onset during spontaneous production. However, any of these top-3 predictions may in fact correspond to the target word itself. This raises a potential confound, since significant decoding performance could be driven by neural activity related to the target word rather than genuine co-activation of multiple lexical alternatives.

To address this possibility, we restricted our analyses to samples where the target word was not among the LLM’s top-3 predictions (Fig. 2A). Here, all three top-ranked predictions are, by definition, unspoken lexical alternatives, which we refer to as the 1st, 2nd, and 3rd alternative respectively from here on (these labels indicate only the LLM-assigned rank and do not assume an equivalent ranking in the human lexical system). If the identities of all three alternatives can be decoded, the effect cannot be attributed solely to neural representation of the produced target and instead provides stronger evidence that multiple lexical alternatives are represented simultaneously. Unless stated otherwise, all subsequent analyses were also limited to samples meeting this criterion.

**Figure 2.**
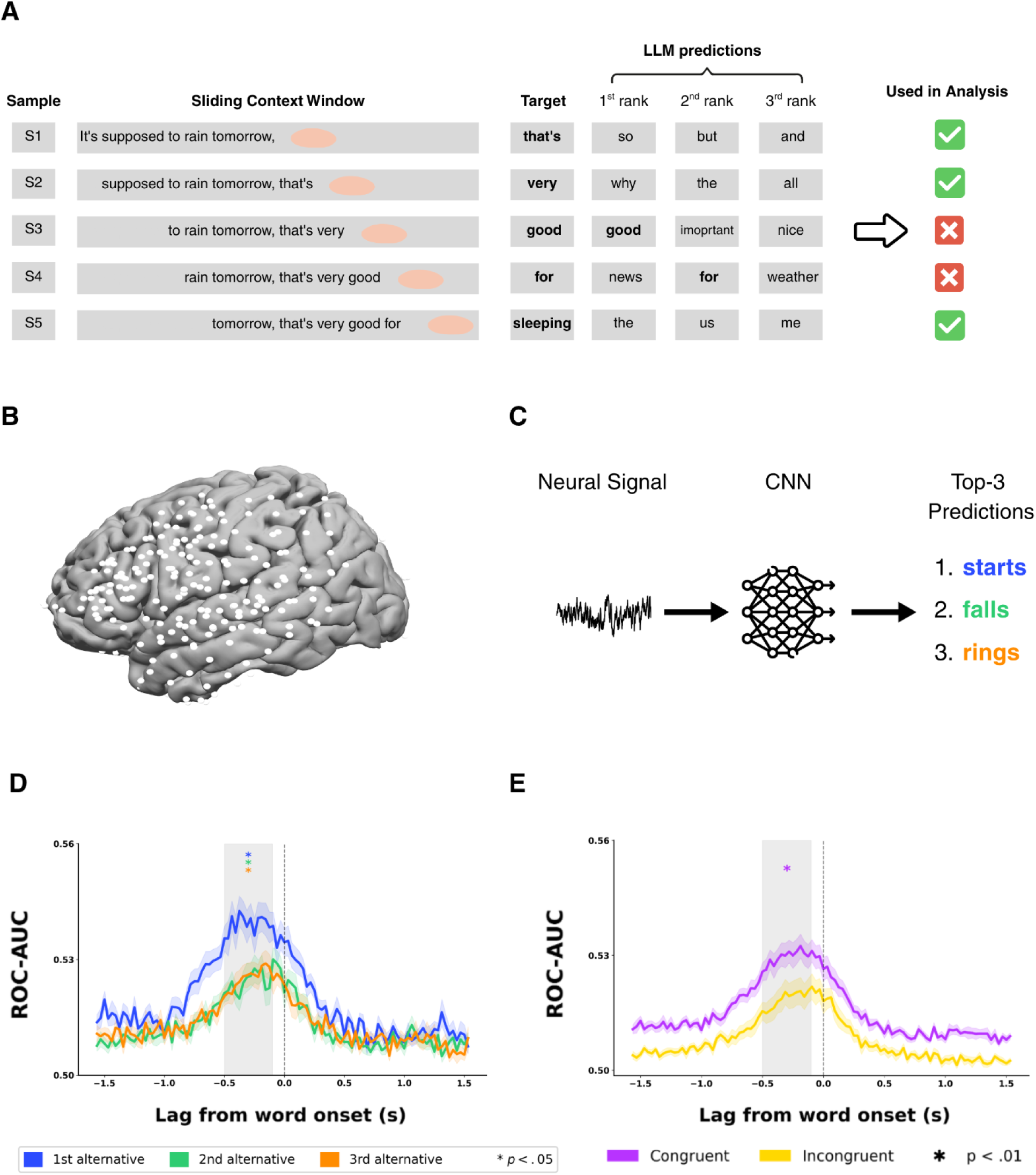
Decoding lexical alternatives from neural activity during production. **A.** Visualization of the sample-filtering procedure: Each row represents one of five samples generated by a sliding context window applied to a single example sentence from the 24/7 dataset. For each sample, the target word (the actual next word in the sentence) and an LLM’s top-3 predicted continuations are shown. To ensure that decoding reflects neural activity related to alternatives (unsaid lexical candidates) rather than the produced word, we only include samples where the target word was not among the model’s top-3 predictions (denoted by ✓). **B**. The electrodes used in the decoding analysis across all patients. **C**. Visualization of the decoding pipeline: Neural signals pooled across the whole brain were used to train three different decoding models using an LLM’s top-3 ranking predictions as labels (1st, 2nd and 3rd alternatives respectively), each one consisting of a two-layer CNN. **D**. In the pre-onset window of interest (−500ms, -100ms) decoding performance (ROC-AUC) was significantly above chance (* p <0.01) for all three alternatives. **E**. To control for similarity between the three lexical alternatives, each model was evaluated on the test data with “congruent” (same labels as training) and “incongruent” labels. For example, a model trained on the 1st alternatives, was tested either on the 2nd alternative labels or 3rd alternatives labels in the incongruent condition. The average performance (ROC-AUC) across models (1st, 2nd and 3rd alternatives) in the “congruent” condition was significantly higher than in the incongruent condition (*p < 0.01). In other words, each alternative can be decoded from the neural activity above and beyond its similarity to the other two lexical alternatives.

Following previous work (Goldstein et al. 2024), we trained a two-layer convolutional neural network (CNN) for each set of labels. Given that only 1 of the 4 patients in the 24/7 dataset had any electrodes implanted in the right hemisphere, we used only left hemisphere electrodes (n = 644). Crucially, three separate decoding models were trained using labels corresponding to the model’s top-3 predictions. Meaning, one decoder was trained to predict the 1st alternative from the neural activity, the other was trained to predict the 2nd alternative from the neural activity, and the third was trained to predict the 3rd alternative from the neural activity.

Each CNN (i.e., decoder) was then evaluated not only on the labels of the alternative it was trained on (“congruent” decoding; e.g., trained on 1st alternative labels and tested on 1st alternative labels), but also on the other two alternative labels (“incongruent” decoding; e.g., trained on 1st alternative labels and tested on 2nd alternative labels, see Fig. 2C). This cross-alternative generalization framework allowed us to directly test whether neural representations associated with different lexical alternatives can be distinguished from another.

Decoding performance, quantified using ROC-AUC, was significantly above chance in the window of interest (−500ms, -100ms) for all top-3 labels (p<0.01, see Methods), demonstrating that the neural signal contains robust predictive information about multiple LLM-predicted alternatives before speech onset (Fig. 2D, see Supplementary for results on all samples). Notably, congruent decoding performance was significantly higher than incongruent decoding performance before onset across participants and across all three alternatives (p<0.01, Fig. 2E). In other words, a decoder trained to identify the 1st alternative generalized significantly better to the 1st alternative than to the 2nd- or 3rd alternatives, and analogous effects were observed for the other alternatives (see Supp. Fig. X for results per alternative).

This dissociation is critical for interpreting the pre-onset neural signal. If decoding performance was driven solely by shared similarity between lexical alternatives, one would expect comparable performance across congruent and incongruent conditions. Instead, the significantly higher performance of congruent decoders indicates that each alternative is represented by a partially distinct neural pattern that can be independently recovered from the population signal. These findings therefore support our interpretation that multiple lexical alternatives for the next word are concurrently represented in the speaker’s brain prior to articulation, and that these representations contain information beyond their mutual similarity.

Taken together, these results show we can separately extract the identity of multiple lexical alternatives (estimated via top-3 LLM predictions) from neural activity before the target word is said. While this end-to-end decoding paradigm allows us to directly probe whether specific lexical alternatives are present in the neural signal, it does leave open questions regarding the nature of these representations. For this reason, we now turn to encoding analyses where we can use different types of embeddings of these alternatives to test how well they capture their representation in neural activity.

#### 1.2 Linear Models Predict Neural Activity from Word Embeddings of Lexical Alternatives

We show that neural activity prior to target word onset can be predicted from semantic embeddings of lexical alternatives, restricting ourselves to the inferior frontal gyrus (IFG), which is known for semantic, syntactic, and to a lesser extent phonological linguistic processing (Sahin et al. 2009; Goldstein, Wang, et al. 2025; Hagoort and Indefrey 2014; Ishkhanyan et al. 2020), and includes Broca’s area. We trained linear encoding models (following (Schrimpf et al. 2021; Goldstein et al. 2022; Antonello et al. 2023; Cai et al. 2025)) to predict neural activity from 50-dimensional semantic word embeddings (GloVe; (Pennington et al. 2014) of these alternatives (see Fig. 1). Because the top-3 predictions represent different alternatives for the same context, we deliberately used static rather than contextual embeddings, so that each alternative could be represented independently. Encoding models were trained independently for each electrode in the IFG at time lags ranging from -2 to +2 seconds relative to word onset (lag 0). As in Section 1.1, we restricted analyses to samples where the top-3 predictions did not include the target word (Fig. 2A), to focus specifically on unsaid lexical alternatives. As a baseline, we also trained an encoding model using semantic embeddings of the target word.

Reproducing results from previous work (e.g., (Goldstein et al. 2022)) we found that the target word significantly predicts neural activity before onset. We also observed significant encoding (see Methods) of the three (unsaid) lexical alternatives using semantic embeddings (Fig. 3A,B), both at the level of individual electrodes (A) and when averaged across the IFG (B). These results provide converging evidence, independent of the decoding analyses above, that multiple lexical alternatives are simultaneously represented in neural activity prior to target articulation during spontaneous production (see Supp. Fig. Y for results per patient).

**Figure 3.**
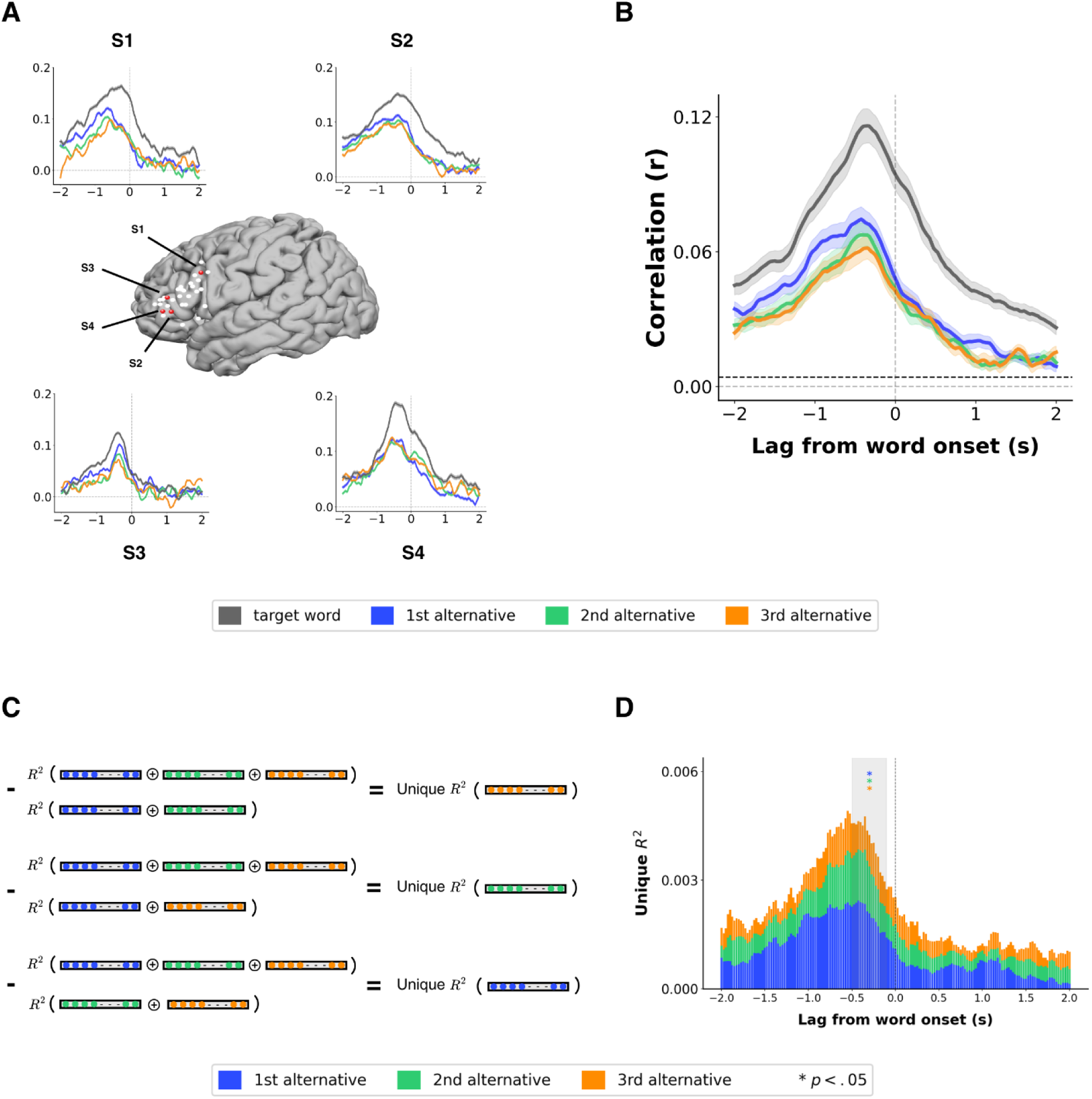
Each lexical alternative captures a unique portion of the neural activity in the IFG during production. **A**. Word-by-word encoding during real-world conversations, shown for four single electrodes (one from each patient) located in the inferior frontal gyrus (IFG). Encoding models were trained on 50-dimensional GloVe embeddings of lexical alternatives, estimated via the top-3 predicted words from GPT-2 XL (blue = 1st alternative, orange = 2nd, green = 3rd, grey= target word). Models were estimated separately for each lag relative to word onset (0 s) and evaluated by computing the correlation between predicted and actual neural activity, with data shown as mean ± s.e. across 10 folds. **B**. Average encoding performance using GloVe embeddings across all electrodes in the IFG, aggregated across all patients. The black dashed line marks the threshold for statistically significant encoding (q < 0.01, two-sided, FWER corrected). **C.** A visualization of the variance partitioning method. To determine how much variance is uniquely explained by each alternative, four new encoding models were fit: a full model containing a concatenation (⊕) of all three lexical alternatives, and three leave-one-out (LOO) encoding models (each containing a concatenation of two out of the three lexical alternatives). The unique contribution of each alternative word was then determined by subtracting the variance explained by the two-way concatenation model without that alternative from the full model. This shows how much variance can be explained by each alternative above and beyond the other two. **D.** Each of the three lexical alternatives explains a unique portion of the variance. The grey shaded area represents the time window of interest between 500ms and 100ms before target word onset, and the asterisks denote that the average unique variance explained by each alternative in this time window is significantly > 0 (p < 0.05).

#### 1.3 Each Lexical Alternative Captures a Unique Portion of the Neural Signal

In the previous analysis, we showed that each of the three lexical alternatives given by an LLM’s top-3 predictions significantly predicts neural activity in the IFG. However, given that these lexical alternatives are often semantically close to each other, this effect could also be a consequence of correlations, or shared information, between them. For example, a lower-ranked alternative might achieve significant encoding simply because of its similarity to that of a higher-ranked alternative (e.g., in Fig. 2, where the 3rd alternative is “better” and the 1st alternative is “good”). To disentangle possible overlaps in information between the lexical alternatives, we use variance partitioning analysis (LeBel et al. 2021; de Heer et al. 2017). This allows us to estimate the unique contribution of each lexical alternative, thus establishing the activation of each of them beyond its similarity to the other two (see Fig. 3C).

To determine how much variance is uniquely explained by each of the lexical alternatives, we first fit a full encoding model with a concatenation of all three lexical alternatives. To estimate the model weights of the model, we use banded ridge regression (Dupré la Tour et al. 2022), which can optimally estimate a different regularization parameter for each of the embeddings concatenated. We then fit three separate encoding models in a leave-one-out (LOO) scheme, each trained on the concatenated embeddings of two out of the three lexical alternatives. To estimate the unique variance explained by each alternative, we then subtracted the variance explained by the corresponding LOO model (for additional details, see Methods) from the variance explained by the full model. This shows how much variance can be explained by each alternative above and beyond the other two, or the similarity between them. A post hoc correction was applied to the estimated variance explained by each model to account for sampling noise and prevent negative values (for more details, see Methods). To see the results without correction, see Supp. Fig. X.

Fig. 3D shows the unique variance in the neural signal in the IFG during production explained by each alternative. Looking at the time window of interest (500ms to 100ms before onset), each of the three lexical alternatives explains a significant portion of the variance (p < 0.05, see Methods), above and beyond the other two. In other words, the variance explained by each alternative cannot be accounted for by the similarity between the lexical alternatives, whether in the embedding space or in the neural activity itself.

Overall, these results provide converging evidence alongside our decoding results that multiple (at least three) lexical alternatives, which can be estimated using an LLM’s top ranked predictions, are represented simultaneously in the speaker’s brain during real-world conversation, each capturing a unique portion of the neural signal. However, there still remains the question of the level of representation of each alternative which we explore in the next section.

#### 1.4 Neural Representation of Lexical Alternatives is Not Just Semantic

Up to this point, we have argued that multiple alternatives are represented in neural activity, while referring to lemmas or lexical items. However, models of word production broadly agree that words can be represented at multiple levels, including semantically (as concepts), syntactically (as lemmas), morphologically, and phonologically. To support our claim that alternatives are indeed represented as lemmas (and not just as semantic concepts), we need to show the neural signal contains syntactic information for each alternative.

To determine which features are represented for each alternative we used the same variance partitioning approach as in Section 1.3 with three different types of embedding representing three different feature spaces: semantic, syntactic and phonological. This allowed us to test whether the syntactic features of each alternative explain variance in the signal beyond that which is explained only by its semantic features. For the semantic feature space we used Glove embeddings as in previous analysis (d=50). For the syntactic feature space, we built a 1-hot embedding representing part of speech (d=50, following (Zada et al. 2025)), and finally for the phonological feature space we built an embedding where each phoneme is represented by 24 articulatory features (d=336, similar to (Goldstein, Ham, et al. 2025), see Methods for more details). We note that Glove embeddings do contain some syntactic information (Pennington et al. 2014), however given that we look at the variance explained by each embedding beyond the other two, this cannot account for any unique variance captured by the syntactic embedding.

For each alternative we fit four encoding models: one full model using the concatenation of all three feature spaces, and three LOO models using the concatenation of only two out of the three feature spaces (see Methods). We then subtracted the variance explained by each of the LOO models from the full model, allowing us to estimate the unique variance captured by each feature space.

Fig. 4 shows the unique contribution of each feature space for each alternative. We see qualitatively similar results between all three alternatives, with the semantic and syntactic feature spaces explaining a significant portion of the variance (p < 0.05, see Methods) in the time window of interest (500ms to 100ms before onset). The phonological feature space, on the other hand, doesn’t explain a significant amount of variance in the specified time window.

**Figure 4.**
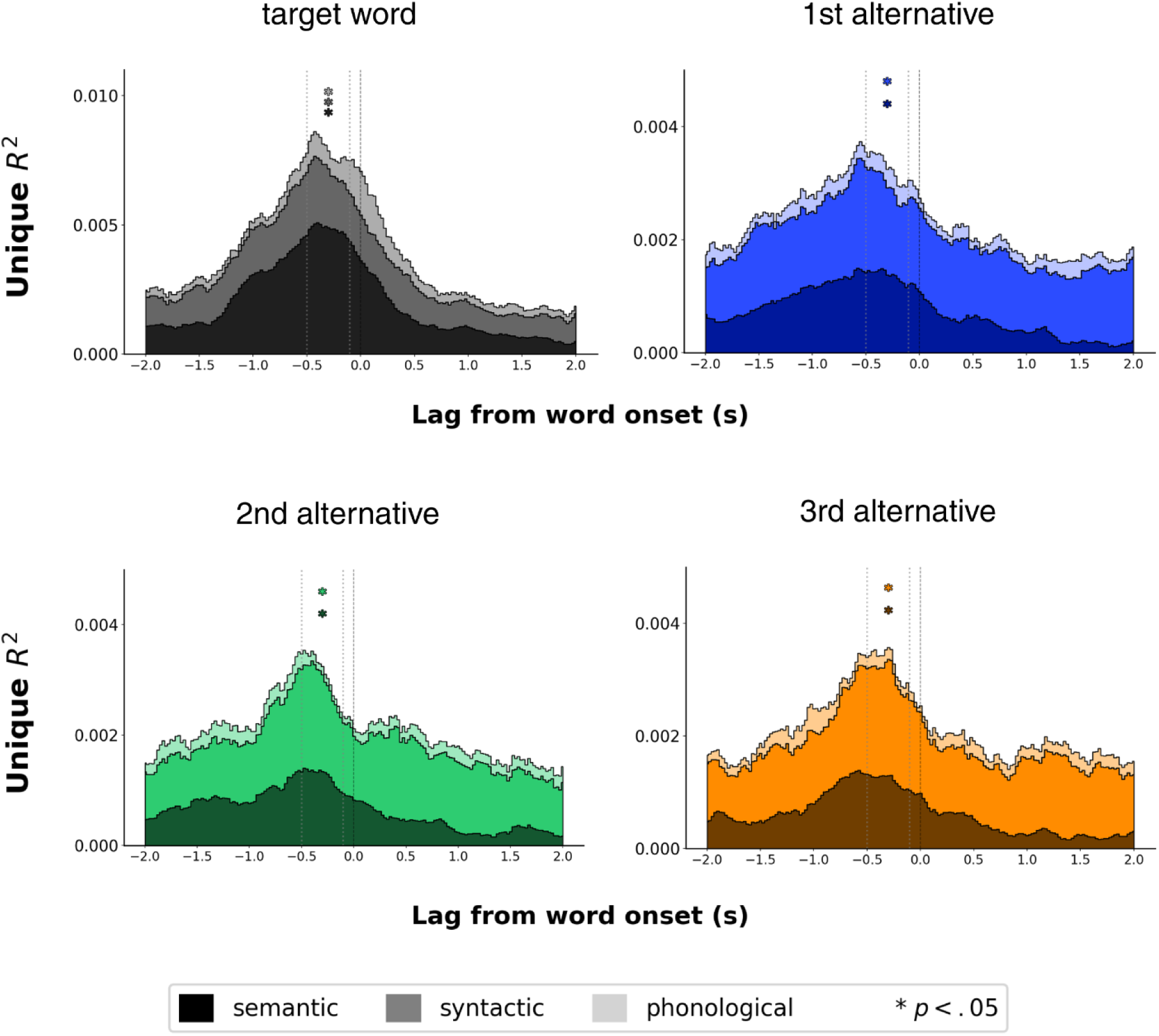
Unique variance explained by each of the LLM’s top-3 predictions (i.e. lexical alternatives). Unique variance explained by each feature space: semantic (d=50), syntactic (d=50) and phonological (d=336) for each alternative and for the target word. For all three alternatives, both the semantic and syntactic features spaces explain unique variance that is significantly > 0 (p < 0.05).

These results support our interpretation that prior to target word onset there are (at least) three distinct unsaid lexical alternatives which are represented in the neural activity as semantic-syntactic entities (i.e. lemmas), and not just as semantic concepts. We leave the question of phonological representation open for future work.

#### 1.5 More than a Dozen Lexical Candidates are Represented Simultaneously in the Brain

Having established that multiple candidates, whose lexical identity can be estimated via top-ranked LLM predictions, are represented in neural activity in real-world production, we can now ask how many candidates are there. In order to estimate the number of lexical candidates, we expanded our encoding analysis using concatenated embedding from Section 1.3 to a larger number of lexical candidates, concatenating embeddings of progressively bigger sets of top-*k* predictions (Fig. 5). To retain sufficient statistical power at higher *k* values, we conducted this analysis using all timepoints in the data, rather than restricting to instances where the target word did not appear among the top-*k* predictions, which would drastically reduce the number of usable samples for each larger value of *k*.

**Figure 5.**
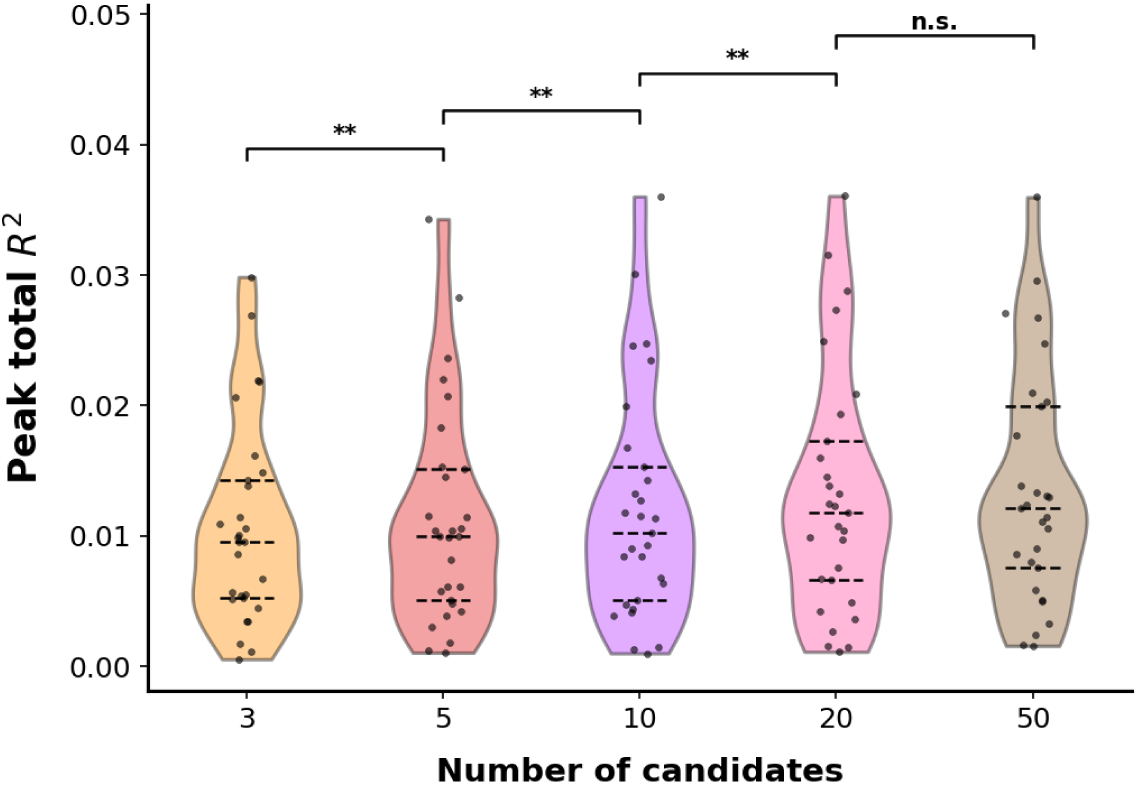
Predicted variance increases with the number of lexical candidates. The peak variance (*R*^2^) explained by encoding models trained on increasing numbers of concatenated (⊕) GloVe embeddings of an LLM’s top-*k* predictions (i.e. lexical candidates) for *k* = 3, 5, 10, 20, 50. The explained variance grew with the number of concatenated candidate embedding, and the highest variance was achieved by the model trained on 50 lexical candidates. Asterisks indicate significant difference in mean variance between consecutive models {(3,5), (5,10), (10,20), (20,50)} (** p < 0.01 with Bonferroni correction for multiple comparisons).

For each *k* ∈ {3, 5, 10, 20, 50} we trained a linear model to predict neural activity from the concatenated embedding of *k* candidates, and then calculated the peak variance (*R*^2^) explained by the model. The peak variance rose monotonically as more lexical candidates were included up to *k*=50, with a significant pairwise difference between the variance explained by *k*=3 and *k*=5, *k*=5 and *k*=10 and *k*=10 and *k*=20 (flip-sign permutation test, p < 0.01 with Bonferroni correction for multiple comparisons, see Methods). The difference between the peak variance explained by concatenating 20 and 50 top ranking predictions was not significant. The surprising fact that using as many as 20 lexical candidates improved the ability of the model to predict neural activity before target onset during production, suggests that a large number of words are represented simultaneously in the speaker’s brain during production prior to articulation.

In summary, we provide, for the first time, multiple pieces of direct neural evidence for the co-activation of multiple lexical alternatives prior to articulation in real-world production, based on decoding and encoding analysis and using top-ranked LLM predictions. In addition, we show that the representations of these alternatives can be disentangled from one another, and that the representation contains both semantic and syntactic information, supporting the claim that these words are represented as distinct lemmas. Expanding this logic from unspoken alternatives to the broader set of lexical candidates, we further show that the amount of neural variance explained continues to grow as more candidates are considered, up to about twenty words, suggesting a surprisingly large candidate set is active in parallel. Next, we explore how the brain updates this set of lexical alternatives during an utterance.

### Principle 2: Lexical Alternatives are Continuously Updated During Conversation On a Word by Word Basis

Given the dynamic and fast paced nature of real-world conversations, we propose that new lexical alternatives are continuously generated for each next word. To test this hypothesis, we split the data into utterances, and then looked at different word positions inside these utterances. Specifically, the first, second and third words in the utterance and the last word in the utterance. This division allowed us to examine multiple consecutive words, as well as take into account the known psycholinguistic effects of the beginning and ending of an utterance, without limiting ourselves to utterances of a certain length. As in previous neural analyses, we used only samples where the top-3 LLM predictions did not include the target word to focus on the alternatives (see Fig. 2A).

We then trained a separate set of encoding models for each word position (i.e., one encoding model trained on concatenated embeddings of LLM predicted top-3 lexical alternatives, and three encoding models trained on concatenated embedding of each pair out of the three lexical alternatives) following the analysis from Section 1.3 to estimate the unique variance explained by each of the top-3 lexical alternatives at this position. Since each word position corresponds only to a small fraction of our data, in order to retain statistical power we used all other word positions as training data and only the focal word position to test the model (e.g., for the first word position, training the models on samples from all other word positions and testing only on first word position samples).

Fig. 6 shows that for all four of the different word positions we tested each of the three lexical alternatives explains a significant portion of the variance (see Methods). These results indicate for each new word in conversation, regardless of its position in the utterance, the brain continuously activates a new set of at least three different lexical alternatives.

**Figure 6.**
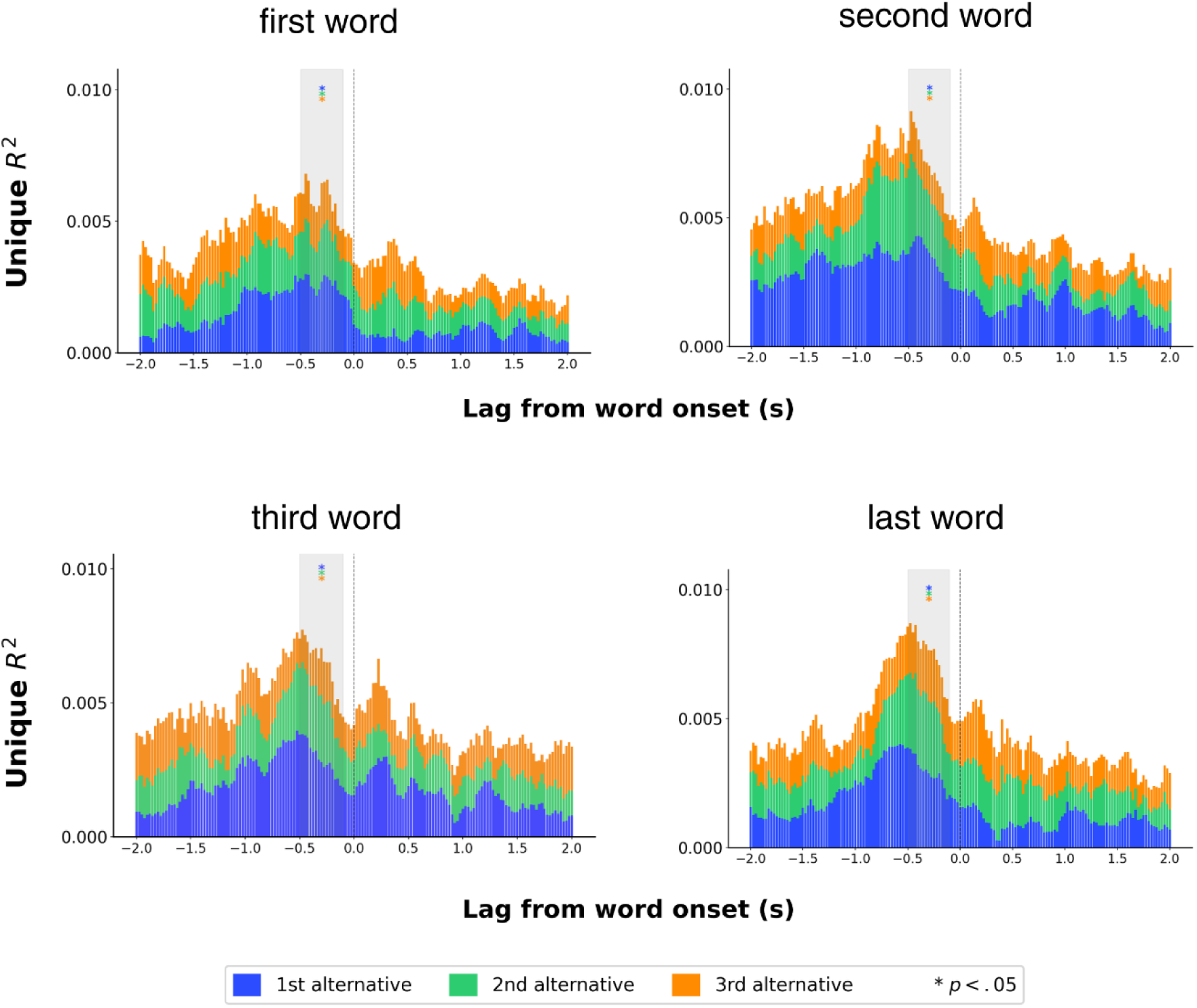
Unique variance predicted by unsaid lexical alternatives at various points in the utterance. A separate set of encoding models was fit for each word position (first, second, third and last words in the utterance), using concatenated (⊕) GloVe embeddings of unsaid lexical alternatives estimated via an LLM’s top-3 predictions. To determine how much variance is uniquely explained by each alternative (at each word position) four encoding models were used: a full model containing a concatenation of all three lexical alternatives, and three leave-one-out (LOO) encoding models each containing a concatenation of two out of three alternatives. The unique contribution of each alternative was then determined by subtracting the variance explained by the LOO model without that alternative from the full model. Asterisks * denote a portion of the variance significantly > 0 (p < 0.05) in the window of interest between 500ms and 100ms before target word onset.

At the same time, as we saw in Section 1.5, there is evidence that a much larger number of words is represented simultaneously, suggesting that perhaps in addition to the lexical alternatives to the current word there are others. Where could these additional words come from? In the next section, we provide evidence showing that these include lexical alternatives to future upcoming words.

### Principle 3: Lexical Alternatives to Multiple Future Words are Represented in the Neural Signal

Up to this point, we have considered the lexical alternatives to the next word for each point in conversation. However, there is evidence that speakers plan a few words ahead (W. J. M. Levelt et al. 1999; Ferreira and Swets 2002), leading us to hypothesize that the brain generates multiple lexical alternatives to each of these planned words as well.

We employed the same paradigm as in Section 1.3 to test whether the lexical alternatives to future target words explain a unique portion of the neural signal. Here, instead of using an LLM’s top-3 predictions for word *n* (given the context up to word *n-1)* to predict the neural signal around the onset of word *n,* we used the top-3 predictions for words *n+1*, *n+2*, *n+3* to predict the neural signal.

As in previous neural analyses, we used only samples where the top-3 predictions represent (unsaid) alternatives (see Fig. 3A) for *n+1*, *n+2*, *n+3*. However, there is still the possibility of overlap between the sets of lexical alternatives of consecutive words. To control for this confound, we further filtered out all samples where these predictions appear in the top-5 predictions for all previous words. For example, if one of the top-3 predictions for *n+3* was either the actual *n+1*th word or appeared in the set of top-3 predictions for *n+2*, this sample was not included in the analysis. This ensures that the encoding achieved by lexical alternatives to target word *n+3* cannot be explained by the representation of lexical alternatives to target word *n+2*.

Fig. 7A shows the variance explained (*R*^2^) by the full models trained on all three concatenated lexical alternatives for *n+1*, *n+2*, *n+3.* All three explain a significant portion of the variance in the window of interest (−500ms, -100ms), suggesting that before the next word is even spoken, multiple lexical alternatives to target words up to three words ahead are represented simultaneously in neural activity.

**Figure 7.**
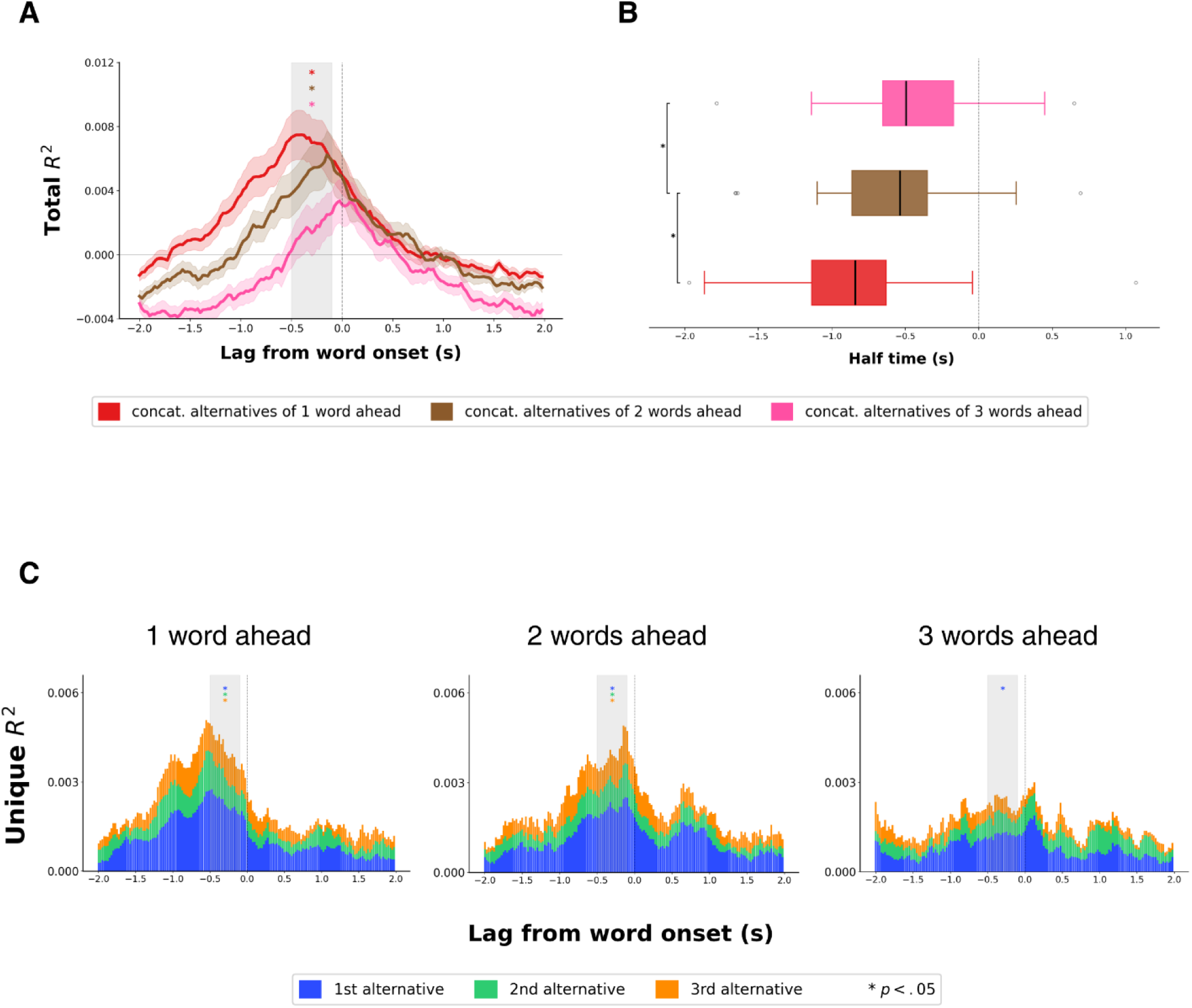
Predicting neural activity using unsaid lexical alternatives to future words. **A.** The total variance (*R*^2^) explained by encoding models trained on concatenated (⊕) GloVe embeddings of an LLM’s top-3 predictions for words *n+1, n+2, n+3* time locked to the word onset of word *n.* For example for *n+2,* in the utterance “I am actually feeling better today”, we would use an LLM’s top-3 predictions for the word “better” to predict neural activity around the onset of the word “actually”. **B**. Box plot of half-times (when the encoding reaches half of the maximum score) of the three models for {1,2,3}-words ahead across electrodes in the IFG. Asterisks, * indicate significant (p < 0.05 with Bonferroni correction) difference between the half times of one and two words ahead, and two and three words ahead. **C**. Unique variance explained by each of the three lexical alternatives to {1,2,3}-words ahead estimated via variance partitioning analysis (see Section 1.3). Asterisks * denote a portion of the variance significantly > 0 (p < 0.05).

Despite the clear temporal overlap between these representations, theoretical accounts of language planning often describe the process as incremental or sequential. To explore the temporal dynamics of future lexical alternatives, we evaluated whether the rise of each set of lexical alternatives happens after that of the previous word. For this, we estimated, for each model, the “half-time” of each neural representation, i.e. the time point where the encoding reaches 50% of its maximal value (following (Zhang et al. 2025)). The difference between the “half-times” of consecutive future words was significant (p < 0.05 with Bonferroni correction, see Methods), for both pairs ((*n+1*, *n+2)*, (*n+2*, *n+3)*). It appears that the brain plans upcoming words in order of their appearance, by first generating lexical alternatives for one word ahead, then two and then three words ahead (Fig. 7A-B).

After establishing that lexical alternatives to multiple words ahead can predict neural activity, we attempted to estimate the unique contribution of each of the three lexical alternatives for each of the future word positions *n+1*, *n+2*, *n+3*. Using the same variance partitioning scheme as in Section 1.4, Fig. 7C shows that for words *n+1*, *n+2* each of the three lexical alternatives explains a significant portion of the variance in the neural signal (beyond the other two), while for *n+3* only the 1st alternative captures variance significantly > 0 (see Methods). We note that this could be due to our conservative method of data filtering described above, which leaves less samples for each consecutive word (since for *each word* we discard samples which overlap with any previous words and lexical alternatives), meaning the *n+3* encoding model has less statistical power.

Taken together, these results suggest that while we speak our brain represents not only lexical alternatives for the upcoming word, but also specific lexical alternatives for multiple words ahead. Furthermore, these sets of lexical alternatives are generated in an incremental fashion according to the order of the corresponding word in the utterance.

## Discussion

The richness of natural language lies in its generative capacity: some thoughts can be verbalized in many different ways. Theoretical models of word production account for this flexibility by positing that word production begins with conceptual preparation of the intended message, which leads to the retrieval of multiple candidates from the mental lexicon, followed by the selection of one of them (Dell 1986; Oppenheim et al. 2010; W. J. M. Levelt et al. 1999). However, previous work has mostly focused on the process of lexical selection, providing only indirect evidence for co-activation in highly constrained setups, leaving the question of neural representation of lexical alternatives in conversational settings largely unexplored. Our use of ECOG data from spontaneous conversations, combined with our novel operationalization of top-ranked LLM predictions, allowed us to estimate a specific set of lexical alternatives to each upcoming word in context. We provide, for the first time, direct evidence for three computational principles governing the organization and temporal dynamics of these alternatives in the brain during real-world production: (1) multiple distinct alternatives are represented in neural activity at the lexical level prior to target articulation; (2) these alternatives are updated continuously on a word-by-word basis; and (3) alternatives to multiple future words are also represented alongside alternatives to the next word.

### Multiple Lexical Alternatives are Represented in Neural Activity in Real-World Production

Using the top-3 ranking LLM predictions as an estimate for a set of lexical candidates at each point in conversation, we showed that the identity of each of these candidates could be decoded from whole brain neural activity prior to articulation. This held true even when looking only at samples where the target (the word that was actually said) was not among the top-3 predictions (Fig 3A,D). The success of decoding models in this setting demonstrates for the first time that human speakers and LLM share *multiple* predictions even when the LLM fails to accurately predict the word that was actually said. This is particularly surprising, given the disagreement between the speaker and the LLM on the identity of the next word. We further found that each of the decoder models performed better on the alternative it was trained on, compared to the other two alternatives (Fig. 3E), suggesting that the representations of the three alternatives in neural activity could be at least partially disentangled from one another.

Turning to encoding analyses, we found that word embeddings of lexical alternatives significantly predicted neural activity in the IFG, and that these representations decayed rapidly, dropping to baseline within one to two seconds (Fig. 3B). This is consistent with prior work suggesting that activation of competing candidates must diminish quickly to support fluent sequential production (Bock and Griffin 2000; Dell 1986). A variance partitioning analysis further showed that each of the three alternatives explained a unique portion of variance beyond the other two (Fig. 3D). Expanding our analysis to larger sets of top-k LLM predictions, we found that the amount of explained variance grew monotonically up to *k* = 20 (Fig. 5), highlighting the surprisingly broad scope of lexical candidates represented in the brain.

Having demonstrated that multiple lexical alternatives were simultaneously and distinctly represented, we examined their representational content using semantic, syntactic, and phonological feature spaces. For all three alternatives, the syntactic features explained a significant portion of the variance beyond the semantic features (Fig. 4). The presence of both semantic and syntactic information strongly suggests that these alternatives are represented as lexical items or at the level of lemmas, and not just as semantic concepts. This is notable given that classic effects such as semantic interference, which were traditionally taken as evidence for lexical co-activation, have alternatively been attributed to pre-lexical conceptual processing, in which multiple concepts are activated without corresponding lexical items (Döring et al. 2022; Kuipers and Heij 2009; Bloem et al. 2004).

Unlike the syntactic features, only the phonological features of the target word (and not the alternatives), were found to explain a significant portion of the variance in the neural signal (Fig. 4). Models of word production typically distinguish lexical retrieval and selection from subsequent phonological encoding (e.g., (Garrett 1975; Kempen 1983)), but disagree on whether these stages overlap. Modular accounts predict that phonological encoding occurs only after target selection, meaning the phonological forms of the alternatives are not active (W. Levelt et al. 1999), whereas cascading and parallel accounts allow phonological activation to spread to multiple lexical candidates (Cutting and Ferreira 1999; Pickering and Strijkers 2025). Although null effects are difficult to interpret, the absence of phonological representation for the alternatives is more consistent with the modular account.

### The Set of Lexical Alternatives is Continuously Updated on a Word by Word Basis

The bulk of prior work concerning lexical retrieval has been focused on the production of single words rather than sentences (Dell et al. 2008), and has mostly utilized picture naming paradigms where the production is in response to static visual imagery. Even in this setting, studies have shown that producing one word changes the accessibility of related words during subsequent retrieval (Howard et al. 2006; Oppenheim et al. 2010), and that interference from distractor words is determined not just by a priori semantic similarity but can be changed on the fly by immediate context creating new connections between words (Abdel Rahman and Melinger 2011). Furthermore, in an interactive picture naming task, it has been shown that lexical retrieval is influenced not only by a speaker’s previous productions but also by words produced by an interlocutor (Hoedemaker et al. 2017). Yet, the question of how lexical alternatives evolve during an utterance in a quick paced conversational setting has remained unexplored.

Looking at different word positions in an utterance (first, second, third and last word), we found that at each one a set of lexical alternatives is represented in neural activity prior to articulation, with each alternative capturing a unique portion of the variance in the signal (Fig. 6). Importantly, the set of alternatives was estimated using the top-ranked predictions of an LLM, which are generated conditioned on the conversational context on a word by word basis. These results show that the identities of neurally represented lexical alternatives are updated at successive word positions in the same utterance, suggesting that conversation continually reshapes the lexical context in which subsequent words are retrieved. Perhaps most surprisingly, multiple alternatives are represented in the brain for the last word in the utterance, even after substantial contextual constraints have accumulated. This implies that even if the speaker knows what they intend to say as they begin to speak, they are still able to reconfigure the wording of their message all throughout the utterance.

### Alternatives to Future Words are Represented Along with Alternatives to the Current Word

Indeed, although many studies over the years have examined planning in language production (Lee et al. 2013; Martin et al. 2014; Swets et al. 2014; Wheeldon et al. 2013), conclusions regarding how many words are actually retrieved ahead of time have been mixed and sometimes contradictory. Some have argued that sentence processing requires advance planning of the syntactic structure of the upcoming phrase, but that lexical retrieval of future words can be delayed until after production onset (Roeser et al. 2019). Others have even suggested that speakers only plan ahead one word at a time (Zhao and Yang 2016). However, to the best of our knowledge, there have been no studies into the retrieval of lexical alternatives to these future target words.

We find that the lexical alternatives of up to three words ahead are represented in neural activity prior to the articulation of the current word, and that for both the next word and the next-next word, each of the three lexical alternatives explains a unique and significant portion of the neural signal (Fig. 7A,C). Furthermore, we also see that the lexical alternatives to future target words are activated in order, so that prior to articulation of target word *i*, first alternatives to target word *i*+1 are activated, then to *i*+2 and then to *i*+3 (Fig. 7A,B). This is in line with theoretical accounts of lexical retrieval, which predict that at the stage where words are represented as lemmas (before phonological encoding), they are activated incrementally, meaning that if in an utterance word *i* precedes word *j*, then word *i* is activated first (e.g., (Dell 1986; MacKay 1982)).

### The Importance of Using Real-World Conversational Data to Study Production

One of our main contributions is the investigation of lexical alternatives in spontaneous conversations. In the past, it has often been assumed that the cognitive and neural mechanisms underlying word production are shared between picture naming and having a conversation (Roelofs 2018). However, naturalistic language studies have shown that conclusions derived from simplified experimental materials do not always fully generalize to real-world language use (e.g., (Hamilton and Huth 2020)). More generally, in recent years, there has been a growing consensus that understanding high level cognition processes requires studying them in ecologically valid settings (Vigliocco et al. 2024; Nastase et al. 2020). In this sense, demonstrating these effects in conversational speech is not merely a replication but an important extension of prior work.

### How are words represented at the computational level?

This question has been at the heart of a long standing debate in cognitive science, with two dominant and seemingly opposing frameworks: *localist* and *distributed* encoding. Localist models represent individual words as discrete nodes embedded within a structured network (W. Levelt et al. 1999; Roelofs 1999; Collins and Loftus 1975), while in distributed models, an individual word is represented by a pattern of activation across many shared units, with meaning arising from the relationships among these patterns (e.g., (Hinton 1986; McClelland and Elman 1986; Elman 1992). Spreading-activation models of word production are typically formulated in localist terms (Dell 1986), where conceptual preparation activates one or more semantic concept nodes, which in turn transmit activation to lexical nodes corresponding to the lexical candidates (which are the focus of this work). Although candidate identity is represented locally in these models, the state of the lexical system at a given moment can be described as a distributed graded pattern of activation across many nodes, implying that both frameworks may be just different views of the same computation. We suggest that LLMs can be used as a framework to capture this dual view: the contextual embedding generated by an LLM, itself a distributed representation of the next word, can be linearly projected onto the vocabulary giving a graded level of activation for each candidate.

### Limitations of LLMs

Despite our findings that top ranked LLM predictions are represented in neural activity in real-world production and their proposed interpretation, we do not suggest that these predictions capture the full set of lexical candidates represented in the brain, nor that these candidates arise through equivalent underlying mechanisms. While recent work has shown close correspondence between the temporal hierarchy of language processing in the brain and the layer-by-layer computation of LLMs (Goldstein, Ham, et al. 2025), other studies have found notable differences in both neural and behavioral responses (Zhou et al. 2023; Vaidya et al. 2023; Wilcox et al. 2021). Together, these results suggest that LLMs may approximate certain aspects of human language processing while diverging in others, an outcome that is perhaps expected given the profound differences in architecture, learning signals, and objectives.

Finally, our study leaves open the question of how one word is ultimately chosen from among the candidates that are co-activated before production. It has long been debated whether this process is competitive or non-competitive in nature (see (Oppenheim and Balatsou 2019) for a review). By providing the first direct neural evidence of parallel candidate activation prior to word production, and by showing that LLM-derived predictions can approximate this latent candidate set, our work lays the groundwork for future investigations into the nature of lexical selection and how it unfolds in the human brain.

## Methods

### Preprocessing the speech recordings

We used a semi-automated preprocessing workflow with four main stages. In the first stage, speech recordings were de-identified by manually removing sensitive information in accordance with HIPAA guidelines. Next, using a human-in-the-loop approach, Mechanical Turk transcribers produced accurate transcripts from noisy, multi-speaker audio. In the third stage, transcripts were aligned to the corresponding audio using the Penn Forced Aligner, with manual corrections applied to ensure precise word-level timing. Finally, speech signals were synchronized with neural data by recording audio through ECoG channels, enabling alignment of neural activity with conversational transcripts at a temporal precision of approximately 20 milliseconds. For a full description of the procedure, see (Goldstein et al. 2023).

### Preprocessing the ECoG recordings

We implemented a semi-automated analysis pipeline to detect and discard corrupted segments of neural data (e.g., periods affected by seizures or electrode instability), and to remove additional noise sources using FFT, ICA, and de-spiking methods (Honey et al. 2012). We then filtered the neural recordings in the high-gamma band (75–200 Hz) and calculated power envelopes as estimates of local neural firing rates (Manning et al. 2009). The resulting signals were standardized via z-scoring, smoothed with a 50 ms Hamming window, and clipped at the boundaries to minimize edge artifacts. All steps were carried out using custom MATLAB 2019a (MathWorks) scripts. For a full description of the procedure, see (Goldstein et al. 2023).

### Prediction and embedding extraction

We extracted contextualized predictions from GPT-2 (gpt2-xl, 48 layers), using the pre-trained version of the model implemented in the Hugging Face environment (Wolf et al. 2019). We first converted the words from the raw transcript (including punctuation and capitalization) to whole or sub-word tokens. We used a sliding window of 1024 tokens (maximum length context of GPT-2 model), moving one token at a time to extract the model’s top-*k* predictions for the final token in the sequence. We then used GloVe (Pennington et al. 2014) word embeddings (d=50) to represent each of these top-k predictions.

### Whole Brain Decoding

To assess whether individual top-ranked candidate words could be independently recovered from neural activity, we performed a decoding analysis using a two-layer convolutional neural network (CNN) with dropout. Neural activity from whole brain electrodes served as input, allowing the model to capture both local and distributed spatial patterns across electrodes. Separate decoders were trained using labels corresponding to the model’s *i*-th ranking predicted next-word candidates (*i*=1,2,3). For each rank condition, decoding performance was evaluated both on the matching rank condition (congruent decoding) and on the remaining rank conditions (incongruent decoding), enabling us to assess the specificity of the decoded representations across candidate ranks.

All decoding models were trained and evaluated using 5-fold cross-validation. Decoding performance was quantified at each temporal lag relative to word onset using the area under the receiver operating characteristic curve (ROC–AUC). To determine whether decoding performance was significantly above chance, we performed a one-sample t-test against chance level (AUC=0.5) independently at each lag across patients and cross-validation folds. Resulting p-values were corrected for multiple comparisons across lags using false discovery rate correction (FDR; Benjamini-Hochberg procedure).

To test whether congruent decoding performance significantly exceeded incongruent decoding performance, we performed a sign-flip permutation test at each temporal lag across patients and cross-validation folds. Specifically, for each patient-fold pair, the sign of the difference between the two decoding conditions was randomly flipped across 5,000 permutations to generate a null distribution of mean differences under the null hypothesis of no systematic performance difference. The observed mean difference was then compared against this null distribution to obtain a two-sided p-value. These p-values were likewise corrected across temporal lags using FDR (Benjamini-Hochberg).

### Electrode-wise encoding

We fit linear regression models for each electrode and temporal lag relative to word onset, predicting neural activity from GloVe embeddings. We defined 161 lags spanning –2,000 to +2,000 ms in 25 ms increments. For each lag, we averaged the neural signal over a 200 ms window, yielding a single value to be predicted from the embedding. To evaluate model performance, we used ten-fold cross-validation: regression weights were estimated with ordinary least squares on the training folds and applied to held-out data. Model accuracy was quantified as the Pearson correlation between predicted and observed neural responses per lag. For a full description of the procedure, see (Goldstein et al. 2022).

### Significance tests

To identify significant electrodes, we implemented a permutation-based randomization procedure. For each iteration, we disrupted the correspondence between embeddings and neural signals by cyclically shifting the word embeddings across time in order to preserve autocorrelation statistics. A different random shift was applied on each permutation, preserving the autocorrelation structure of the neural data while eliminating any true alignment between words and brain activity. Importantly, we used GloVe embeddings of the target word in this procedure to ensure that there was no overlap with model-derived alternatives. For every electrode, we repeated the full encoding analysis 5,000 times. In each iteration, we retained the maximum encoding value across all lags, and then, to control for multiple comparisons across electrodes, we further retained the maximum of these values across all electrodes. This yielded a null distribution of 5,000 maximum values. For each electrode, we then computed a P-value as the percentile rank of its observed (non-permuted) maximum encoding value relative to this null distribution. Finally, we applied false discovery rate (FDR) correction (Benjamini and Hochberg 1995), considering electrodes with q < 0.01 to be significant.

To determine a threshold for significant average encoding values (black line in Fig. 3, 4 and 5), we implemented a permutation-based procedure with 1000 permutations per electrode. Using embeddings of the 3rd ranking predictions in the “all” setting, we reran the encoding analysis for each permutation and computed the average encoding value across all electrodes within a given region of interest (IFG). For each lag, we derived a null distribution of average encoding values and estimated a threshold corresponding to α = 0.01. To correct for multiple comparisons across lags, we adopted a max-statistic approach: specifically, we retained the maximum threshold across all lags, thereby controlling the family-wise error rate (FWER; (Nichols and Hayasaka 2003)). Observed average encoding values exceeding this corrected threshold were considered significant.

Following Goldstein et al. (2022), we tested the significance of lag-wise differences between two encoding models applied to the same group of electrodes using a sign flip permutation test. For each electrode, we obtained encoding values from both models and then randomly swapped their assignments across models. Repeating this procedure 5,000 times, we computed the average pairwise difference at each lag to generate a null distribution. We then compared the observed differences against this distribution to obtain P-values, which were corrected for multiple comparisons across lags using false discovery rate (FDR) adjustment. Lags were considered significant at a threshold of q < 0.01.

### Unique Variance Analysis

To quantify the variance in neural activity uniquely explained by each lexical alternative, we performed a variance-partitioning analysis (adapted from (LeBel et al. 2021)). For each patient and electrode, we fitted a full encoding model using the concatenated GloVe embeddings of the first, second, and third lexical alternatives, as well as three reduced models in a leave-one-out (LOO) scheme, each containing two of the three embeddings. Models were fitted using banded ridge regression, which assigns a separate regularization parameter to each alternative (Dupré la Tour et al. 2022).

Denoting the three alternative embeddings as A, B, and C, the following nested models were fit:

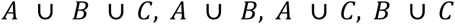

The unique variance explained by each alternative beyond the other two, or relative complement (RC), was calculated as the variance explained by the full model minus that explained by the corresponding LOO model without that alternative:

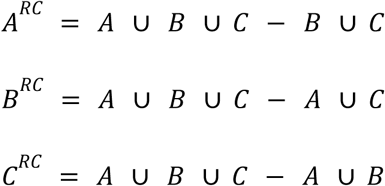

Because sampling noise can cause a reduced model to appear to explain more variance than its corresponding full model, producing theoretically invalid negative unique-variance estimates, we applied the post hoc bias-correction procedure introduced (following (de Heer et al. 2017), (Dupré la Tour et al. 2022)). This procedure estimates the smallest adjustment to the model-performance values required to satisfy the constraint that all unique-variance partitions are nonnegative.

Statistical significance of the unique variance was assessed using 1,000 permutations, with the complete variance-partitioning and bias-correction procedure repeated separately for each permutation, patient, and electrode. For each alternative, corrected unique-variance estimates were averaged across electrodes and time points within the prespecified lexical-retrieval window of -500 to -100 ms relative to target-word onset and then averaged across patients. The observed group-level value was compared with the corresponding permutation-derived null distribution, using a significance threshold of (p<0.01), a conservative threshold that also accounts for the three alternative-specific comparisons according to a Bonferroni correction.

#### Linguistic Feature Spaces Variance Partitioning

To determine which linguistic features of each lexical alternative were represented in neural activity, we performed a feature-space variance-partitioning analysis separately for the first, second, and third alternatives. Each alternative was represented using three feature spaces capturing semantic, syntactic, and phonological information.

Semantic information was represented using 50-dimensional pretrained GloVe word embeddings. Syntactic information was represented using a 50-dimensional one-hot vector indicating the alternative’s part-of-speech tag. Fine-grained part-of-speech tags were assigned to each word in isolation using spaCy (Honnibal et al. 2020), rather than in the original sentence context, because an unselected alternative was not necessarily syntactically compatible with that context. To construct the phonological feature space, each word was first converted into a phoneme sequence using the Phonemizer Python package (Bernard and Titeux 2021). Each phoneme was then represented using articulatory features obtained with PanPhon (Mortensen et al. 2016). Phoneme sequences were zero-padded to 14 positions, corresponding to the maximum number of phonemes in a word in our dataset. Using 24 features per phoneme, this procedure yielded a 336-dimensional phonological representation for each word (14*24 = 336).

For each lexical alternative, we fitted a full encoding model using the concatenation of its semantic, syntactic, and phonological feature spaces. We additionally fitted three reduced models in a leave-one-feature-space-out scheme, each containing two of the three feature spaces. Models were fitted using banded ridge regression, with a separate regularization parameter estimated for each feature space. The unique variance explained by a given feature space was calculated by subtracting the variance explained by the corresponding reduced model from that explained by the full model.

Thus, for example, the unique contribution of the syntactic feature space reflected variance explained by the alternative’s part-of-speech representation beyond that explained by its semantic and phonological representations. The same post hoc bias-correction and permutation-testing procedures described for the lexical-alternative variance-partitioning analysis were applied here.

## Acknowledgements

We are grateful to Timna Wharton Kleinman, Zaid Zada, Noam Siegelman and Mariano Schain for their helpful feedback and valuable insights, and to Miriam Havin for her assistance with figure design.

This work was supported by the European Research Council (ERC) under the European Union’s Horizon Europe research and innovation programme (ERC Starting Grant “WordOrigions”, Grant Agreement No. 101222215).

